# Hydrogen Peroxide induces resistance to DNA damage in a localization and p53 dependent manner

**DOI:** 10.64898/2026.05.13.724825

**Authors:** Janneke P. Keijer, Paulien E. Polderman, Wytze T.F. den Toom, Paula Sobrevals-Alcaraz, Robert van Es, Diego Montiel González, R. Kok, Soufiane El Baghdadi, M. Can Gülersönmez, Edwin. C. A. Stigter, Harmjan R. Vos, Boudewijn M. T. Burgering, Ruben van Boxtel, Tobias B. Dansen

## Abstract

Organisms need to be able to adapt to a changing environment in order to survive. The adaptive response invoked by a low dose of a stressor resulting in resistance to high levels of that stressor is known as *hormesis* and can even lead to lifespan extension of organisms. The exact mechanisms underlying stress-induced hormesis are unknown, although multiple studies pose mitochondria-derived Reactive Oxygen Species (ROS, e.g. H_2_O_2_) as an important contributor. Here we used chemo-genetic H_2_O_2_ production as a model to study ROS-dependent adaptive responses in a localization-dependent manner. We found that brief, sublethal H_2_O_2_ production at the nucleosomes provides p53-dependent resistance to a subsequent high dose of H_2_O_2_, whereas mitochondrial H_2_O_2_ production, surprisingly, does not. A multi-omics approach revealed that p53-induced hormesis is accompanied by metabolic rewiring that boosts reductive capacity, and that the increased stress resistance can mostly be attributed to its downstream target p21. Importantly, brief p53 stabilization also mounted protection against chemotherapy-induced DNA damage, suggesting that p53-dependent hormesis could be exploited to selectively protect healthy, p53-wildtype tissue from chemotherapy in the treatment of patients with p53 mutant tumors.

## Introduction

The ability to adapt to environmental changes and stress is a fundamental property of living systems, reflecting the evolutionary principle that organisms capable of enduring and adjusting to adversity are most likely to persist. This fundamental concept plays a key role in processes at multiple levels of organization in biology including tissue homeostasis, training through physical exercise^1^, lifespan extension^2^ as well as therapy resistance in cancer treatment^3^. At the cellular level, stressors trigger an adaptive response making cells less sensitive to subsequent exposures. Hormesis is a form of adaptation that is characterized by a biphasic, concentration-dependent response to a stressor: low levels of certain stressors lead to an adaptive response and increased resistance to stress that can even extend lifespan in model organisms, whereas higher levels of the same stressors overwhelm the adaptive response, induce toxicity and shorten lifespan^4,5^. This phenomenon is often paraphrased in popular culture with (an adaptation of) the famous quote by Friedrich Nietzsche^6^: “What doesn’t kill you makes you stronger”. The response to Reactive Oxygen Species (ROS) displays a similar biphasic, concentration-dependent response. Whereas supraphysiological levels of ROS can lead to oxidative distress, damage and even cell and organismal death^7,8^, redox signaling^9^ by low levels of ROS are thought to play a role in the adaptive response that protects cells and organisms from subsequent damage^10–15^. Differences in response to ROS are highly dependent on the type of ROS, the subcellular site of production and (local) concentration^7^. Moreover, it has been suggested that in some cases enhanced mitochondria-derived ROS triggers the adaptive response responsible for the hormetic effect through redox signaling^9,16^ and the term *mitohormesis* was coined^17–19^. The rationale underlying this hypothesis is that a stress-induced increase in mitochondrial ROS production activates signaling, resulting in the transcription of genes that promote stress resistance, resulting in an increase in health and lifespan^19,20^.

The cytotoxicity of many chemotherapies have been shown to stem in part from elevated ROS levels, although often the type of ROS and underlying mechanisms are unknown^21^. On the other hand, Mitochondria-derived ROS have also been suggested to contribute to resistance of cancer cells to chemotherapy^22,23^. For example, adaptation to high levels of chemotherapy-induced ROS by elevating antioxidant levels makes cancer cells chemoresistant^24,25^. Similarly, mitochondrial ROS were suggested to drive chemotherapy resistance in non–small cell lung cancer cells^26^. Likewise, inducing mitochondrial oxidative stress by knockdown of SOD2 in mice and mouse embryonic fibroblasts elevates expression of antioxidants and improved resistance to oxidative stress^27^. However, evidence for the direct involvement of (mitochondrial) ROS in hormesis remains circumstantial and is often based on the loss of ROS-induced hormesis upon treatment with antioxidants^13^. Moreover, redox signaling and oxidative stress have often been studied using exogenous addition of H_2_O_2_ or other means of chemical perturbation of the cellular redox state, and it has been questioned how well these methods mimic (patho) physiological redox signaling in terms of concentration, localization and temporal dynamics^28^. The use of ectopic expression of D-amino acid oxidase (DAAO) from *R. gracilis* at different subcellular locations made it possible to study effects of near-physiological H_2_O_2_ levels in a localized manner^29^. Using this system, we previously showed that only nuclear ROS is mutagenic and that exposure to a bolus of exogenous H_2_O_2_ is not mimicking physiological effects^30^. Addition of D-amino acids such as D-Alanine (D-Ala) allows for the inducible and titratable production of H_2_O_2_ by localized DAAO. Using this model system, we here show that low amounts of H_2_O_2_ can induce a hormetic response, and that this depends on where it is produced in the cell (in this study we define hormesis as an adaptive response that renders cells more resistant to a subsequent, otherwise damaging or toxic insult). We found that DAAO-dependent H_2_O_2_ production indeed boosted resistance to a variety of subsequent (genotoxic) stressors. Strikingly, the hormetic response was only observed when DAAO was localized to the nucleosomes, whereas mitochondria-localized DAAO failed to induce hormesis. We set out to delineate the molecular pathways involved in localized H_2_O_2_ dependent hormesis using a multi-omics approach, and pinpointed pathways downstream of p53 as important factors. Stabilization of p53 not only protected against DNA damage and cell death induced by H_2_O_2_, but also prevented accumulation of DNA-damage by chemotherapy. Our observations shed light on therapy resistance development in p53 wildtype tumors, and provide a framework for future strategies to reduce the collateral toxicity of cancer therapeutics in the treatment of p53 mutant tumors.

## Results

### Nuclear H_2_O_2_ induces hormesis, whereas mitochondrial H_2_O_2_ release does not

To study whether ROS can induce hormesis and whether this is dependent on its site of production, we used ectopic expression of D-amino acid oxidase (DAAO) to locally produce H_2_O_2_ in human RPE1-hTERT cells, which are untranformed, have a relatively stable karyotype and a p53 wild-type response to stress. We expressed DAAO at the nucleosomes (H2B-DAAO), to mimic H_2_O_2_ production close to the DNA, and at the cytosolic side of the outer mitochondrial membrane (TOM20-DAAO), to mimic mitochondrial H_2_O_2_ release (Fig. 1A). Intermembrane space (IMS) and mitochondrial matrix (MLS) targeted DAAO were used to model intramitochondrial ROS production. Monoclonal cell lines were selected based on comparable amounts of H_2_O_2_ production upon addition of D-Alanine^30^. Although mitochondrial ROS is normally produced in the form of •O_2_^−^, SOD1 and SOD2 rapidly convert this to H_2_O_2_, the latter ROS is the second messenger in redox signaling.

**Fig. 1:**
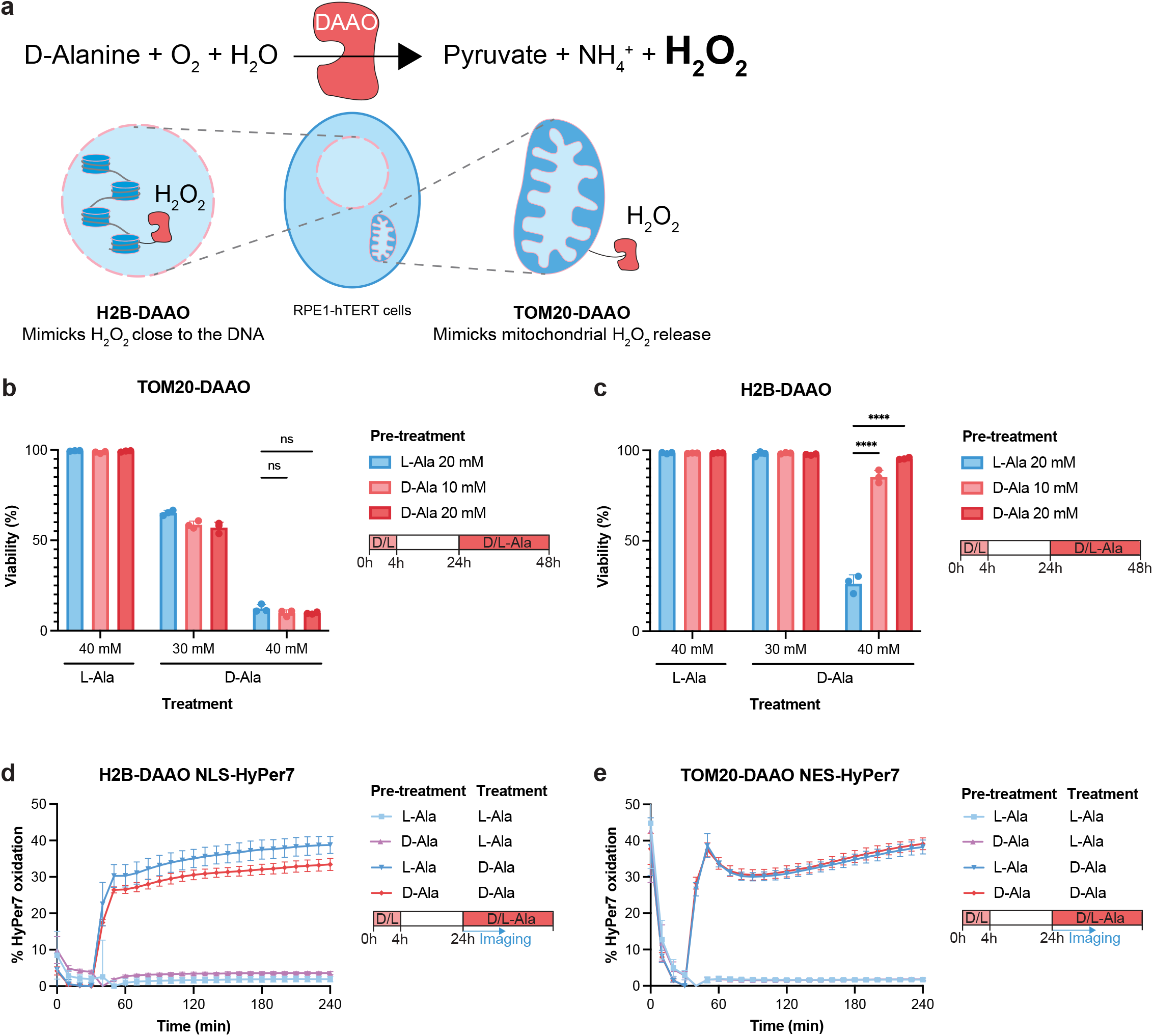
Nuclear H_2_O_2_ induces hormesis, whereas mitochondrial H_2_O_2_ does not. **a** Schematic representation of RPE1-hTERT cells expressing H2B-DAAO or TOM20-DAAO to locally produce H_2_O_2_ upon D-alanine treatment. **b-c** Quantification of cell viability by PI exclusion of RPE1-hTERT TOM20-DAAO cells **(b)** and RPE1-hTERT H2B-DAAO cells **(c)** that were pretreated with L-Ala or D-Ala for 4 hours, followed by 20 hours of recovery and treatment with L-Ala or D-Ala. Dots represent 3 biological replicates of ∼10.000 cells each. Data represents mean +/- SD. Two-way Anova with Bonferroni correction was performed (ns p > 0.05, ****p≤ 0.0001). **d-e** HyPer7 measurements of RPE1-hTERT H2B-DAAO cells with nuclear NLS-HyPer7 **(d)** and RPE1-hTERT TOM20-DAAO cells with cytosolic NES-HyPer7 **(e)** that were pretreated with 10 mM L-Ala or D-Ala for 4 hours followed by 20 hours of recovery. Subsequently, cells were imaged and 30 mM L-Ala or D-Ala was added. Maximal HyPer7 oxidation after 400 uM H_2_O_2_ treatment was set as 100% and basal HyPer7 ratio was set to 0%. Data represents mean +/- SD of 4 biological replicates. Treatment was started after the HyPer7 baseline was stable following transfer to the microscope (30 minutes).

To test whether a short pulse of H_2_O_2_ produced at the mitochondria or in the nucleus would protect cells to a subsequent, otherwise lethal dose, we pretreated cells with a low, sublethal dose of D-Ala for 4 hours, followed by 20 hours of recovery. We then assessed whether this affected survival in response to a 24-hour treatment with a high dose of DAAO-dependent H_2_O_2_ production. Low amounts of H_2_O_2_ production at the outer mitochondrial membrane did not affect survival during a high dose of H_2_O_2_ produced at the outer mitochondrial membrane (Fig. 1B) or exposure to exogenous H_2_O_2_ generated by Glucose Oxidase in the media (Supplementary Fig. 1C). Likewise, intramitochondrial H_2_O_2_ did not induce hormesis, as neither H_2_O_2_ production at the inter-membrane space nor in the mitochondrial matrix increased resistance to a following treatment (Supplementary Fig. 1A-B). Conversely, sublethal concentrations of H_2_O_2_ produced at the nucleosomes resulted in a marked increase in resistance to a subsequent high dose of nuclear H_2_O_2_ production (Fig. 1C) or extracellular H_2_O_2_ (Supplementary Fig. 1D). A 4-hour pretreatment with D-Ala did not induce observable cell death, excluding the possibility that the observed acquired resistance to H_2_O_2_ stems from the selection of intrinsically more resistant cells. This suggests that a low dose of nuclear, but not mitochondrial H_2_O_2_ induces a hormesis-like adaptive response in our model.

### Nuclear H_2_O_2_ increases reductive capacity of the cell

The observed increased resistance to H_2_O_2_ following brief H2B-DAAO activation could be due to a higher cellular reductive capacity, which could facilitate more efficient scavenging of H_2_O_2_ through the peroxiredoxin/thioredoxin/thioredoxin reductase and glutathione peroxidase/glutathione reductase systems, both of which depend ultimately on NADPH as electron donor. We performed live fluorescence microscopy using HyPer7, a ratiometric, genetically encoded fluorescent H_2_O_2_ sensor that reports on the combined rate of oxidation (by H_2_O_2_) and reduction (by the Trx system)^31^. We show that pretreatment with a low dose of nuclear H_2_O_2_ production, followed by 20 hours of recovery resulted in lower oxidation levels of HyPer7 in the nucleus (NLS-HyPer7) in response to a subsequent treatment with a high dose of nuclear H_2_O_2_ production (Fig. 1D). A lower HyPer7 oxidation/reduction ratio could in theory also be caused by a decrease in H_2_O_2_ production by DAAO upon hormesis induction. To exclude this, we assessed H_2_O_2_ production based on oxygen consumption rates, using a method described before^32^. H_2_O_2_ production by DAAO was not affected by pretreatment with nuclear H_2_O_2_ (Supplementary Fig. 1E), indicating that the observed decrease in HyPer7 oxidation/reduction ratio can be attributed to an increased reductive capacity. Unlike H2B-DAAO, low level TOM20-DAAO activation followed by recovery did not affect the reductive capacity as measured by NES-HyPer7 upon subsequent TOM20-DAAO activation, (Fig. 1E), corroborating the absence of adaptation and resistance to a high dose of H_2_O_2_.

Besides a decrease in HyPer7 oxidation levels, we also observed a decrease in the oxidation/reduction ratio of the fluorescent GRX1-roGFP sensor following brief H2B-DAAO activation (Supplementary Fig. 1F), indicating that the ratio of oxidized versus reduced glutathione (GSSG/2GSH ratio) is shifted towards more reduced glutathione. In conclusion, these results demonstrate that production of sublethal amounts of H_2_O_2_ at the nucleosomes, but not at the mitochondria, induces an adaptive response that increases the reductive capacity of the cell. Because both the Trx and GSH-dependent systems are affected, an increased rate of NADPH regeneration could potentially be at the basis of this observation.

### Nuclear H_2_O_2_ induced hormesis is associated with the upregulation of p53-related pathways

To determine the adaptations that are responsible for inducing hormesis upon nuclear H_2_O_2-_ induction, we looked at differences in gene expression after 4 hours of sublethal H2B-DAAO activation and compared this to TOM20-DAAO activation, which did not induce hormesis in our system. Transcripts from p53-related genes like MDM2, CDKN1A and TIGAR were induced when H_2_O_2_ was produced at the nucleosomes, but not at the outer mitochondrial membrane (Fig. 2A-B). Gene set enrichment analysis indeed identified that the p53 pathway was upregulated exclusively in H2B-DAAO (Fig. 2C, Supplementary Fig. 2A-B). Genes associated with the hallmark pathway Reactive Oxygen Species^33^ were also enriched upon H_2_O_2_ treatment in H2B-DAAO (NES = 1.78, padj = 0.00851; Supplementary Fig. 2B) but not in TOM20-DAAO, although this pathway was not among the top 10 enriched. mRNA sequencing data after 4 hours of D-Ala gave an insight into the direct transcriptional responses to local H_2_O_2_ production, but the hormetic effect in Fig. 1 was assessed 20 hours after removal of the low dose of D-Ala. By this time, the initial transcriptional response to H_2_O_2_ production likely has ceased. We therefore also quantified the changes in the proteome following 4 hours D-Ala plus 20 hours recovery, to identify proteins and processes that may underlie nuclear H_2_O_2_-induced hormesis. In line with the results from the mRNA-Seq experiment, label-free quantitative proteomics identified p53-pathway related proteins expressed at higher levels after induction of nuclear H_2_O_2_ (TIGAR, CDKN1A etc.) (Fig. 2D), but not upon mitochondrial H_2_O_2_ release (Supplementary Fig. 2C). Protein set enrichment analysis indeed revealed upregulation of p53 pathways (Supplementary Fig. 2D) after brief exposure to sublethal nuclear H_2_O_2_, posing p53 as a candidate involved in hormesis induction.

**Fig. 2:**
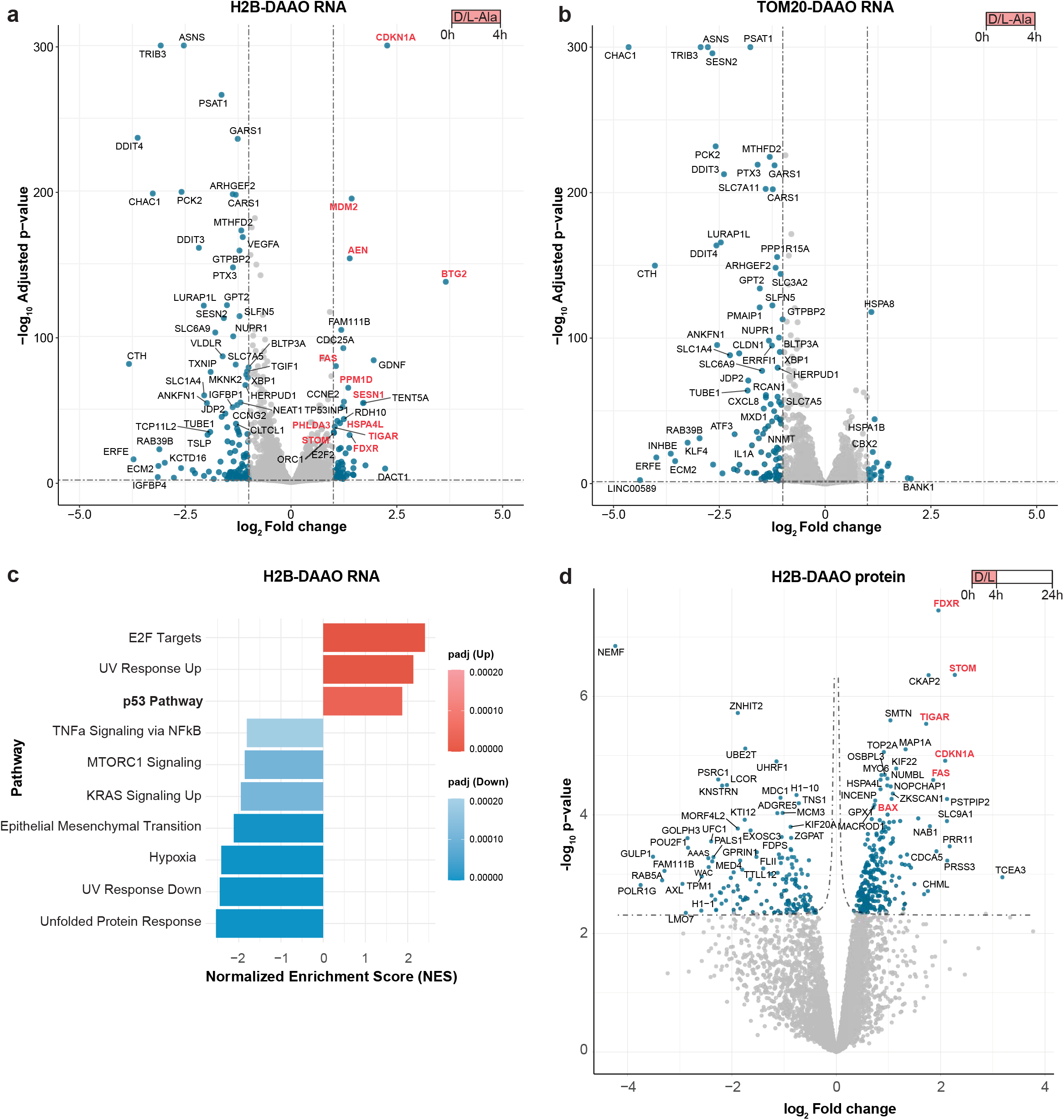
Nuclear H_2_O_2_ induced hormesis is associated with the upregulation of p53-related pathways. **a-b** Volcano plot depicting differentially expressed genes assessed by mRNA sequencing comparing L-Ala versus D-Ala treatment (10 mM) for 4 hours in RPE1-hTERT H2B-DAAO cells **(a)** and RPE1-hTERT TOM20-DAAO cells **(b)**. Dots on the right represent genes with higher expression levels in cells treated with D-Ala and dots on the left represent genes with higher expression levels in cells treated with L-Ala. Genes related to p53 are highlighted in red. Blue dots represent significantly upregulated or downregulated genes (log_2_ Fold change >1 or <-1 and -log_10_ Adjusted p-value <0.05). Data represents 3 biological replicates per condition. **c** Top 10 Hallmark pathways that are changed upon D-Ala treatment of RPE1-hTERT H2B-DAAO cells, determined using gene set enrichment analysis (GSEA) of RNA sequencing data of RPE1-hTERT H2B-DAAO cells. **d** Volcano plot depicting differentially expressed proteins of mass spectrometry data comparing L-Ala versus D-Ala treatment of 20 mM for 4 hours, followed by 20 hours of recovery in RPE1-hTERT H2B-DAAO cells. Blue dots represent significantly upregulated or downregulated proteins (FDR≤0.05). Data represents 3 biological replicates per condition.

### Brief stabilization of p53 is sufficient to induce hormesis

In accordance with the observations of the enrichment analyses, brief stabilization of p53, by a 4 hour treatment with MDM2-inhibitor Nutlin-3a, was sufficient to protect against cell death induced by nuclear DAAO-dependent H_2_O_2_ production initiated after 20 hours recovery (Fig. 3A), and the effect was further enhanced when cells were treated with Nutlin-3a for 24 hours. All cells survived in the control (L-Ala) second treatment condition, meaning that Nutlin-3a pretreatment did not kill the cells, again arguing against selection of resistant clones. Additionally, a BrdU incorporation assay showed that (at least in our model system) cells can fully resume cycling after recovering from a brief Nutlin-3a treatment (Supplementary Fig. 3A-B). Surprisingly, Nutlin-3a pretreatment did not protect cells against cell death caused by H_2_O_2_ produced by DAAO at the outer mitochondrial membrane (Fig. 3B) and only minimally to intramitochondrial H_2_O_2_ (Supplementary Fig. 3C). We wondered whether 24 hours of H_2_O_2_ production at or in the mitochondria could overwhelm a p53-depedent hormetic effect and thus obscure protection at earlier time points. To test this, we performed live imaging with propidium iodide (PI) in the culture medium to visualize the accumulation of dead cells over time (PI accumulates on the DNA of dead cells upon loss of membrane integrity). Also in this experiment p53-stabilization protects cells against subsequent nuclear H_2_O_2_ production, also at earlier time points (Supplementary Fig. 3D), but not against mitochondrial H_2_O_2_ production (Supplementary Fig. 3E).

**Fig. 3:**
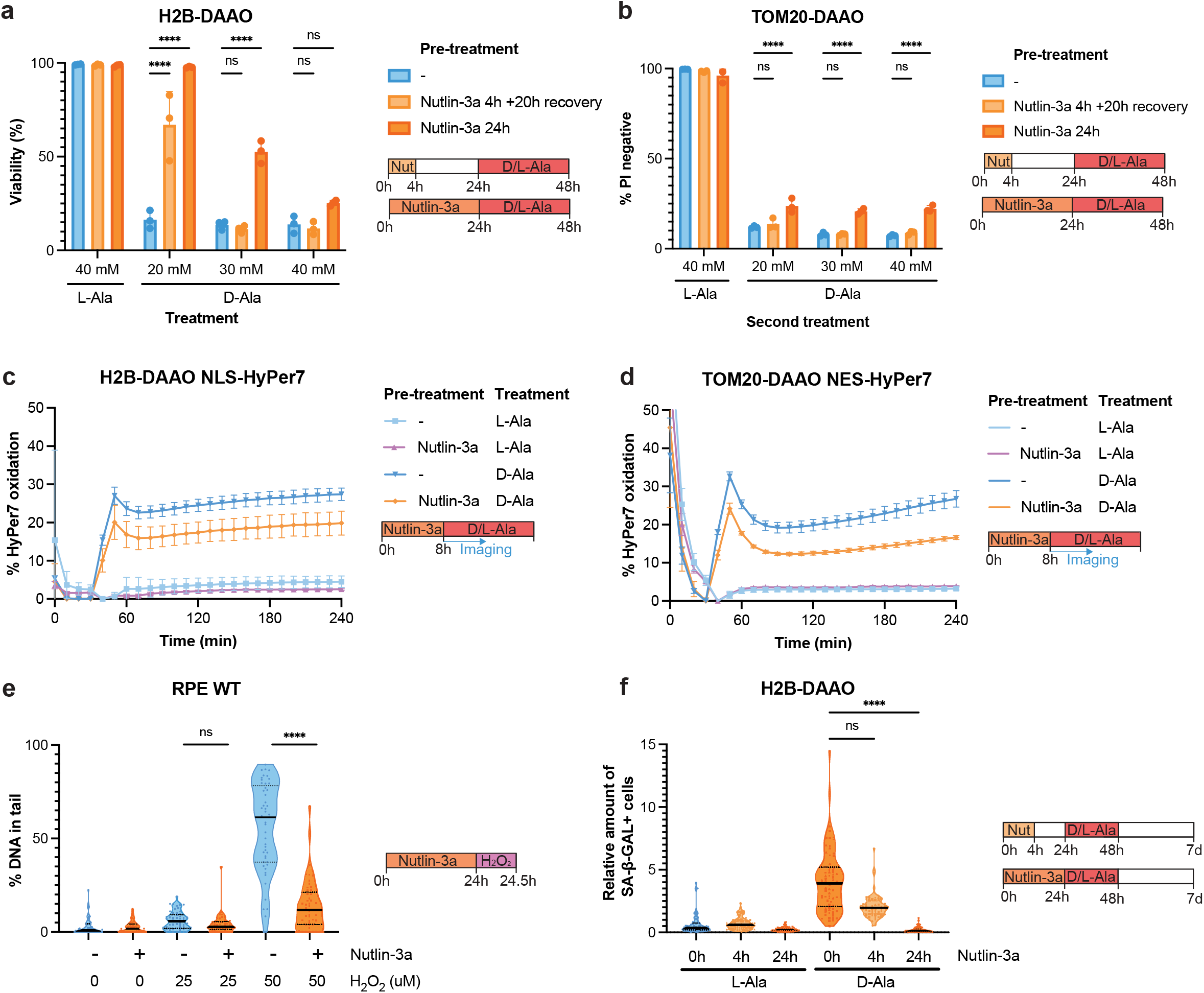
Brief stabilization of p53 is sufficient to induce hormesis. **a-b** Quantification of cell viability by PI exclusion of RPE1-hTERT H2B-DAAO cells **(a)** and RPE1-hTERT TOM20-DAAO cells **(b)** that were pretreated with Nutlin-3a (10 uM) for 4 hours and 20 hours of recovery or for 24 hours, followed by treatment with L-Ala or D-Ala. Dots represent 3 biological replicates of ∼10.000 cells each. Data represents mean +/- SD. Two-way Anova with Bonferroni correction was performed (ns p > 0.05,****p≤ 0.0001). **c-d** HyPer7 measurements of RPE1-hTERT H2B-DAAO cells with nuclear NLS-HyPer7 **(c)** and RPE1-hTERT TOM20-DAAO cells with cytosolic NES-HyPer7 **(d)** that were pretreated with Nutlin-3a (10 uM) for 8 hours. Subsequently, cells were imaged and 30 mM L-Ala or D-Ala was added. Data represents mean +/- SD of 4 biological replicates. **e** Quantification of alkaline comet assay of RPE1-hTERT WT cells pretreated with Nutlin-3a (10 uM) for 24 hours, followed by 30 min of H_2_O_2_ treatment. Line represents mean of ∼50 comets per condition. Kruskal-Wallis test with Dunn’s post-hoc test was performed (ns p > 0.05, ****p≤ 0.0001). **f** Quantification of Senescence associated-β-galactosidase (SA-β-gal) staining of RPE1-hTERT H2B-DAAO cells that were pretreated with Nutlin-3a (10 uM) for 4 hours and 20 hours of recovery or for 24 hours, followed by treatment with 10 mM L-Ala or D-Ala for 48 hours. Subsequently, cells were kept in culture for 4 more days to allow the SA-β-gal positive senescence phenotype to develop. Representative images are shown in Supplementary Fig. 3I. Dots represent measurements of 3 biological replicates. Kruskal-Wallis test with Dunn’s post-hoc test was performed (ns p > 0.05, ****p≤ 0.0001).

We hypothesized that resistance to nuclear but not mitochondria-derived H_2_O_2_ production that we observed upon p53 stabilization could be explained by a selective increase in reductive capacity in the nucleus. Nevertheless, stabilization of p53 resulted in lower nuclear as well as cytosolic HyPer7 oxidation levels in response to H2B-DAAO- or TOM20-DAAO-dependent H_2_O_2_ production respectively (Fig. 3C-D). In response to exogenous addition of H_2_O_2_, the effect of the p53-dependent increase in reductive capacity was only observed in the nucleus (Supplementary Fig. 3F-G). H_2_O_2_ production by DAAO was not affected by Nutlin-3a pretreatment (Supplementary Fig. 3H), indicating that the lower HyPer7 oxidation levels can be attributed to an increased reductive capacity. This observation suggests that although stabilization of p53 increases reductive capacity throughout the cell, it fails to protect from cell death induced by high amounts of H_2_O_2_ at the mitochondria. p53 must therefore protect against nuclear H_2_O_2_ via another or additional mechanism than through increasing reductive capacity. Although both H2B-DAAO- or TOM20-DAAO-dependent H_2_O_2_ production can induce cell death, only nuclear H_2_O_2_ production leads to DNA damage (as previously shown in ^30^). We thus hypothesized that p53 stabilization selectively protects against (nuclear H_2_O_2_-induced) DNA damage. In line with this, pretreatment of cells with Nutlin-3a for 24 hours protected from H_2_O_2_-induced DNA breaks, as measured by the alkaline comet assay (Fig. 3E). This suggests that p53 stabilization boosts survival by lowering the DNA damage that accumulates upon exposure to H_2_O_2_. We have previously shown that prolonged production of nuclear H_2_O_2_ can induce senescence^30^. In line with the observed protection from DNA damage, Nutlin-3a pretreatment of RPE1-hTERT-H2B-DAAO cells prevented senescence induction upon D-Ala treatment as measured by the senescence-associated-β -gal assay (Fig. 3F, Supplementary Fig. 3I).

### p53-induced hormesis is accompanied by metabolic rewiring

One of the downstream targets of p53 that was upregulated upon nuclear H_2_O_2_ treatment in both RNA sequencing data (Fig. 2A), as well as proteomics data (Fig. 2D), is TP53-induced glycolysis and apoptosis regulator (TIGAR). We confirmed upregulation of TIGAR upon nuclear H_2_O_2_ production by Western Blot (Supplementary Fig. 4A).

TIGAR lowers cellular Fru-2,6-P2 levels, thereby inhibiting glycolysis while promoting the pentose phosphate pathway (PPP), which enhances NADPH regeneration and consequently lowers ROS levels^34^. We hypothesized that TIGAR could be involved in p53-induced hormesis, by increasing reductive capacity of the cell. Metabolomics analysis indeed showed a decrease in glycolysis intermediates (Fig. 4A) and no change or an increase in PPP intermediates (Fig. 4B) upon 24h Nutlin-3a treatment. Nutlin-3a treatment also shifted the GSH/GSSG balance towards more reduced glutathione (Fig. 4C), which is in accordance with the GRX1-roGFP measurements (Supplementary Fig. 1F). Removing glucose from the culture media, decreases the ability of the cell to produce NADPH via the PPP. In line with this, the Nutlin-3a-induced increase in reductive capacity was indeed abolished by removing glucose (Fig. 4D). In absence of glucose, cells become more sensitive to DAAO-derived H_2_O_2_ (Fig. 4E-F). However, Nutlin-3a still was still capable of inducing hormesis when glucose was removed (Fig. 4F). This suggests that Nutlin-3a does increase reductive capacity by increasing the glucose flux to PPP and that this could contribute to hormesis, although another part of the effect is independent of glucose.

**Fig. 4:**
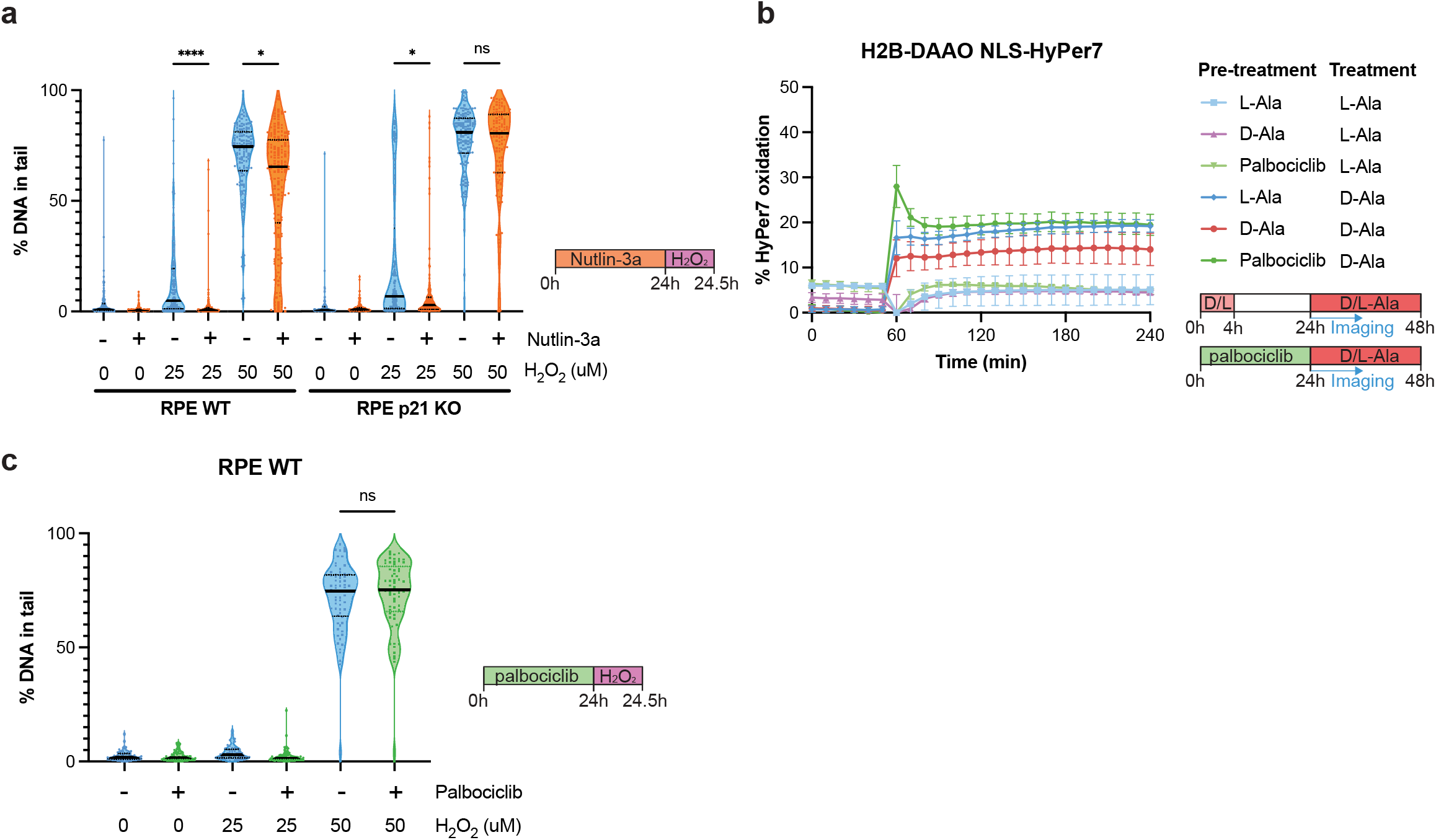
p53-induced hormesis is accompanied by metabolic rewiring. **a-c** Fold change in metabolites measured upon 24h Nutlin-3a treatment compared to untreated cells. Samples were normalized to protein concentration as measured by a BCA assay and subsequently normalized to untreated control. Unpaired t-test with Welch correction and Benjami-Hochberg FDR correction was performed. Data represents 4 biological replicates per condition. **d** HyPer7 measurements of RPE1-hTERT H2B-DAAO cells with nuclear NLS-HyPer7 that were pretreated with Nutlin-3a (10 uM) for 8 hours. Subsequently medium was replaced with medium without glucose, cells were imaged and 30 mM L-Ala or D-Ala was added. Data represents mean +/- SD of 4 biological replicates. **e-f** Quantification of cell viability by PI exclusion of RPE1-hTERT H2B-DAAO cells that were pretreated with Nutlin-3a (10 uM) for 24 hours, medium was replaced with medium with glucose **(e)** or without glucose **(f)** and L-Ala or D-Ala treatment was added. Dots represent 3 biological replicates of ∼10.000 cells each. Data represents mean +/- SD. Two-way Anova with Bonferroni correction was performed (ns p > 0.05, *p≤ 0.05, **p≤ 0.01, ***p≤ 0.001, ****p≤ 0.0001).

### p53-induced hormesis is largely mediated by p21, but not by cell cycle arrest alone

p53 activation results in the transcriptional induction of a multitude of downstream effects, of which p21-dependent cell cycle arrest arguably is the most canonical. Based on our transcriptomic and proteomic data (Fig. 2A, Fig 2D), we hypothesized that p21-dependent cell cycle arrest upon p53 activation could be a driver of the observed hormetic effects. We therefore investigated p53-induced hormesis in p21 knock-out cells, in which activation of p53 does not result in a cell cycle arrest (Supplementary Fig. 5A). In the absence of p21, p53 activation by Nutlin-3a no longer protected to H_2_O_2_-induced DNA damage at a high dose of H_2_O_2_ and only showed a minor effect at a lower dose of H_2_O_2_ (Fig. 5A). This suggests that p53-induced hormesis can be mainly attributed to p21.

**Fig. 5:**
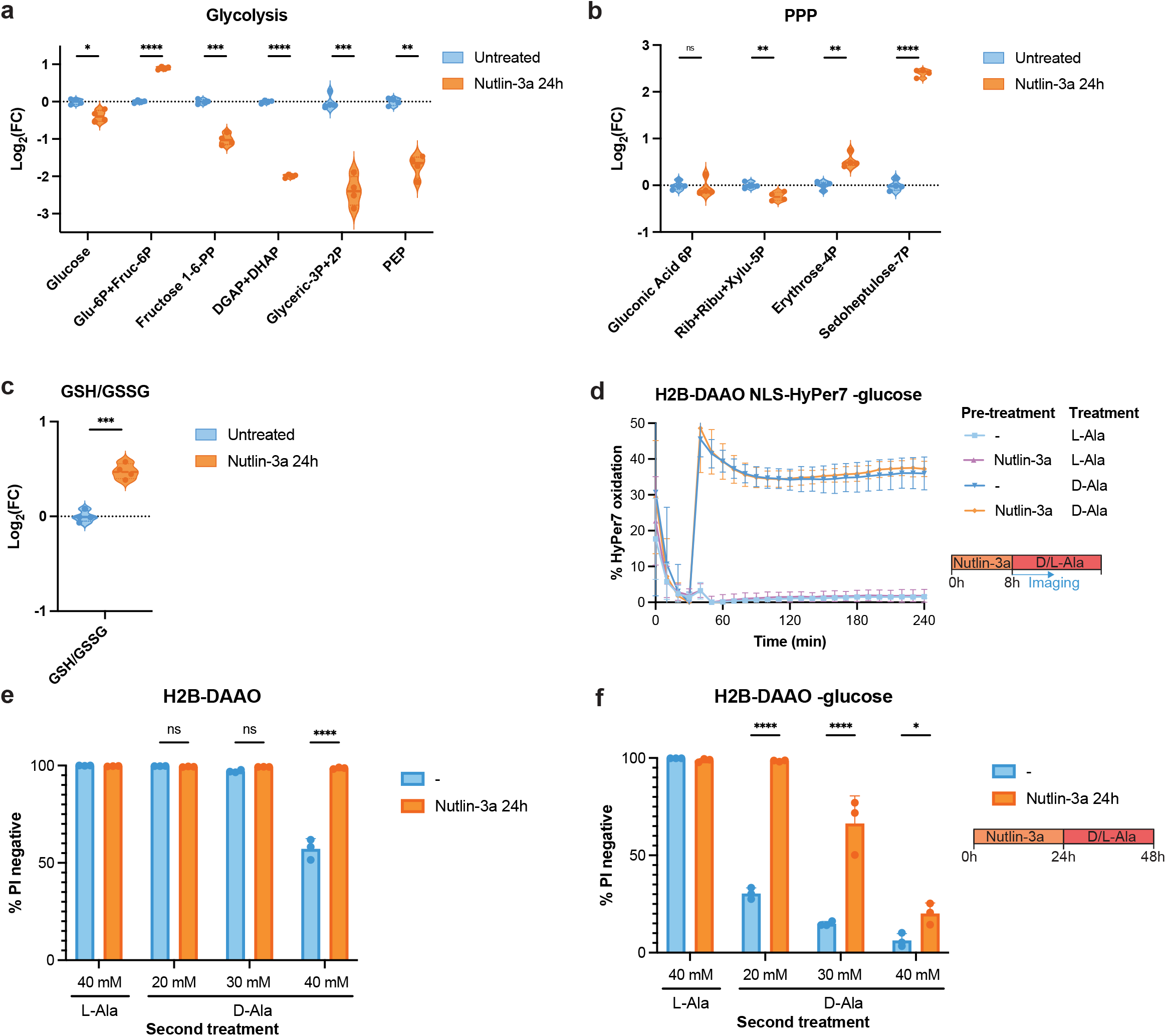
p53-induced hormesis is largely mediated by p21, but not by cell cycle arrest alone. **a** Quantification of alkaline comet assay of RPE1-hTERT WT cells and RPE1-hTERT p21 KO cells pretreated with Nutlin-3a (10 uM) for 24 hours, followed by 30 min of H_2_O_2_ treatment. Line represents mean of ∼50 comets per condition. Kruskal-Wallis test with Dunn’s post-hoc test was performed (ns p > 0.05, *p≤ 0.05, ****p≤ 0.0001). **b** HyPer7 measurements of H2B-DAAO cells with nuclear NLS-HyPer7 that were pretreated with 20 mM L-Ala or D-Ala for 4 hours and 20 hours of recovery or palbociclib (250 nM) for 24 hours. Subsequently, cells were imaged and 25 mM L-Ala or D-Ala was added. Data represents mean +/- SD of 4 biological replicates. **c** Quantification of alkaline comet assay of RPE1-hTERT WT cells pretreated with palbociclib for 24 hours, followed by 30 min of H_2_O_2_ treatment. Line represents mean of ∼50 comets per condition. Kruskal-Wallis test with Dunn’s post-hoc test was performed (ns p > 0.05).

We used the CDK4/6 inhibitor palbociclib to test whether hormesis is also induced when the cell cycle is blocked at the G1/S border by other means. Palbociclib pretreatment did not lead to a decrease in oxidation levels upon H2B-DAAO-dependent nuclear H_2_O_2_ production (Fig. 5B) and did not result in lower amounts of H_2_O_2_-induced DNA damage (Fig. 5D). Because it has been suggested that palbociclib can in some cases also activate p53^35^, we performed this experiment also in p53 KO cells and confirmed the observation that cell cycle arrest by palbociclib pretreatment has no effect on protection from DNA damage (Supplementary Fig. 5B). The efficient induction of cell cycle arrest in both p53 wild-type and p53 KO cells by palbociclib treatment was confirmed by flow cytometry using cells treated in parallel with the HyPer7 imaging and comet assays (Supplementary Fig. 5C). In conclusion, a cell cycle arrest by itself does not lead to lower H_2_O_2_-induced DNA damage induction, whereas p21 induction upon p53 stabilization does. We deduce that the p53-induced hormetic effect extends beyond the induction of cell cycle arrest.

### p53-dependent hormesis in the context of chemotherapy

Having shown that stabilization of p53 can protect cells to future oxidative stress-induced DNA damage and cell death, we wondered whether this adaptive response extends to resistance to other DNA-damage inducing treatments commonly used in the clinic as anti-cancer therapeutics. Indeed, Nutlin-3a pretreatment protected to DNA damage caused by Etoposide (Fig. 6A) or Doxorubicin (Fig. 6B) in cells that are wild type for p53, while leaving p53 KO cells vulnerable. We even observed a slight increased vulnerability of p53 KO cells to Etoposide (Fig. 6A). Interestingly, p53 stabilization did not protect against gamma-irradiation-induced DNA damage (Fig. 6C).

**Fig. 6:**
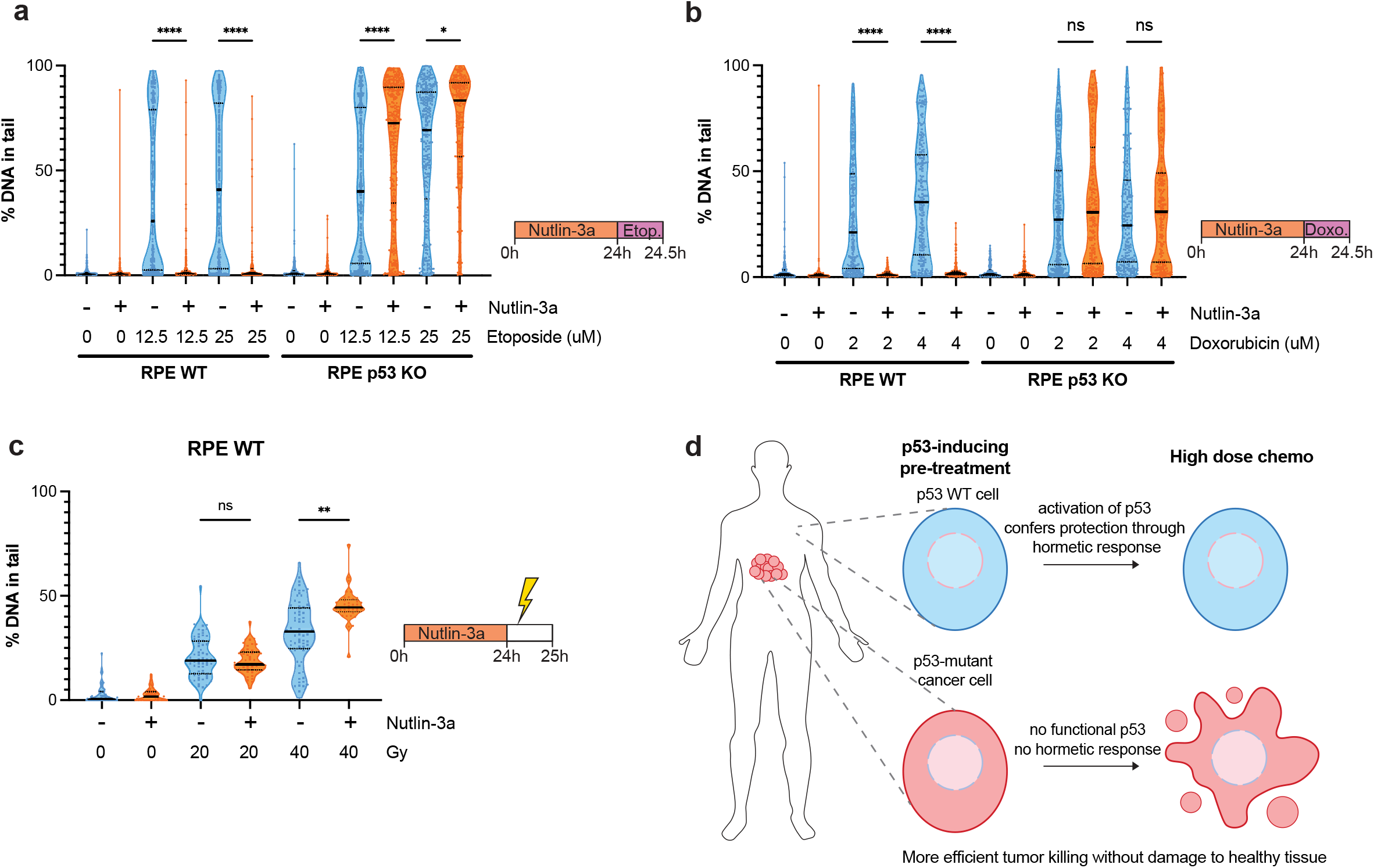
p53-dependent hormesis in the context of chemotherapy. **a-b** Quantification of alkaline comet assay of RPE1-hTERT WT cells and RPE1-hTERT p53 KO cells pretreated with Nutlin-3a (10 uM) for 24 hours, followed by 30 min of etoposide **(a)** or doxorubicin **(b)** treatment. Line represents mean of ∼130 comets per condition. Kruskal-Wallis test with Dunn’s post-hoc test was performed (ns p > 0.05, *p≤ 0.05, ****p≤ 0.0001). **c** Quantification of alkaline comet assay of RPE1-hTERT WT cells pretreated with Nutlin-3a (10 uM) for 24 hours, followed by irradiation. Cells were harvested 1 hour after irradiation. Line represents mean of ∼50 comets per condition. Kruskal-Wallis test with Dunn’s post-hoc test was performed (ns p > 0.05, **p≤ 0.01). **d** Schematic representation of potential therapeutic application. Patients with a tumor that is mutant for p53 might benefit from a p53-inducing pretreatment (such as Nutlin-3a). This would induce p53-dependent hormesis selectively in healthy tissue, thereby protecting from chemotherapy-induced collateral damage and side-effects. The p53 mutant tumor on the other hand does not induce hormesis and remains sensitive to the treatment.

These observations could have translational implications to be tested in future work: a p53-dependent hormetic response could in theory boost resistance of (p53 wild type) healthy tissue to collateral DNA damage from chemotherapy, while leaving the p53 mutant tumor tissue vulnerable (Fig. 6D). A downside of such a strategy could be that the timing and dose of Nutlin-3a treatment must be carefully chosen, which is not trivial in patients. Prolonged p53 stabilization could provoke a chronic cell cycle arrest or apoptosis, thereby hampering function, especially in proliferative tissue like the bone marrow or the intestinal epithelium. In our model system, we showed that cells can fully resume cycling after recovering from a brief Nutlin-3a treatment (Supplementary Fig. 3A-B). Taken together, our data indicate that there could be a therapeutic window to exploit p53-dependent hormesis to protect healthy tissue from collateral DNA damage from chemotherapy in the context of tumors that are devoid of p53 function.

## Discussion

Studies in both cultured cells and model organisms have suggested a role for ROS in triggering an adaptive response to stress that can serve to drive resistance to future stresses^11–15^. Recent work has led to the notion that ROS (mainly in the form of H_2_O_2_) is highly compartmentalized, and that phenotypes downstream of perturbations of redox homeostasis depend on where H_2_O_2_ is produced^7^. We therefore investigated whether a ROS-dependent adaptive response can be triggered in a chemogenetic model for localized H_2_O_2_-production (RPE1-hTert-DAAO). Our work suggests that such a ROS-dependent adaptive response can indeed be triggered by localized H_2_O_2_, but only when this is produced close to the DNA. Although mitochondria are a major source of ROS, and ROS-induced hormesis has often been attributed to mitochondrial ROS production, we found no adaptative response to H_2_O_2_-production inside or at the mitochondria in our model system. Low-level H2B-DAAO activation followed by recovery protected to subsequent, and otherwise lethal, H_2_O_2_-produced at the nucleosome (by H2B-DAAO) or extracellularly (by Glucose Oxidase in the media). We showed that this nuclear H_2_O_2_-induced hormetic effect is caused by activation of p53 and that activation of p53 alone by Nutlin-3a could also induce protection against H_2_O_2_. In line with this observation, low dose radiation-induced hormesis that was described in mice was also shown to be dependent on p53^5^. It remains to be established whether nuclear H_2_O_2_ and p53 activation is also involved in other models for hormesis, but unlike what we found in human cells, a recent study showed that low level mitochondrial matrix localized DAAO activation triggered stress resistance in the fission yeast *S. Pombe*^36^. Endogenous sources of enzymatic nuclear H_2_O_2_ production include Lysine-specific histone demethylase 1 (LSD1)^37^ and NADPH oxidase NOX4 and MICAL1 have also been shown to contribute to nuclear ROS levels^38–40^, but it remains to be established whether and under what conditions these sources could mount a similar adaptive response. In the context of cancer treatment, many (DNA-damage-inducing) chemotherapies have been shown to be accompanied by elevated nuclear ROS levels (reviewed in ^41^). Our data suggest that this chemotherapy-induced nuclear ROS may trigger an adaptive response, which could contribute to therapy resistance in p53 wild type tumors. Whereas p53-dependent hormesis upon low-level H2B-DAAO activation in our study coincides with DNA damage, p53 stabilization can also occur for instance upon prolonged treatment with the thiol-specific oxidant diamide, which induces more oxidizing conditions in both the nucleus and the cytosol, but in the absence of DNA damage^42^. TOM20-DAAO activation to produce H_2_O_2_ at the cytosolic face of mitochondria did not induce nuclear ROS nor DNA damage and did not result in p53 stabilization (Fig 2 and ^30^). We propose that oxidizing conditions in the nucleus, for instance due to elevated H_2_O_2_, activates a p53-dependent hormetic response irrespective of DNA damage. In line with this, treatment with the MDM2 inhibitor Nutlin-3a, which stabilizes p53 also in the absence of DNA damage, efficiently induced a similar hormesis phenotype. Our -omics data did not indicate a role for NRF2 in the protective effect, although this transcription factor is often implicated in the upregulation of antioxidant capacity in response to ROS^43^.

Mechanistically, we showed that p53-induced hormesis was largely dependent on its transcriptional target p21, which is required for induction of a cell cycle arrest in G1 phase downstream of p53 activation (see e.g. Supplementary Fig. 5A). There could be various explanations for why a cell cycle arrest would protect against DNA damage. Different DNA damage repair pathways are active in each cell cycle phase^44^, so it could be that damage does occur, but is more quickly repaired when cells are arrested downstream of p53 activation. This scenario seems unlikely, as we do not see protection against DNA breaks induced by irradiation (Fig. 6C). The Nutlin-3a induced cell cycle arrest even seems to sensitize cells to irradiation, which is in line with previous research showing that a G2 arrest sensitizes to irradiation^45^. Alternatively, cell cycle progression might be needed for acquiring DNA breaks during DNA replication. For example, oxidative base lesions can cause double-strand breaks by causing replication stress^46,47^. Likewise, Etoposide induces DNA breaks by trapping topoisomerase II on DNA loops, which results in a road block and prevents topoisomerase from relaxing DNA supercoiling^48^. Similarly, doxorubicin acts by intercalating in the DNA and trapping topoisomerase II onto the DNA, blocking replication and the transcriptional machinery, but also by generating ROS^49^. Induction of a cell cycle arrest could therefore in principle mitigate the damaging effects of these chemotherapies. Similarly, it has been described before that antagonists of MDM2 can protect p53 wild-type cancer cells from DNA damage induced by mitotic inhibitors^50,51^, and that this depends on p21-dependent cell cycle arrest. However, H_2_O_2_ not only induces oxidative base lesions, but also directly damages the DNA backbone^52^, which is likely irrespective of DNA replication. Furthermore, if a (G1) cell cycle arrest would fully explain the DNA damage resistance phenotype, one would expect that treatment with the CDK4/6 inhibitor Palbociclib would exert the same effect, which was not the case (Fig. 5C). We therefore conclude that p21 either has a functional role besides cell cycle arrest in the observed hormetic response, or that protection against DNA damage requires the interplay of p21-dependent cell cycle arrest and another p53-regulated transcriptional program.

The observed increase in reductive capacity that accompanies the hormetic response could be due to activation of the p53 transcriptional target TIGAR. TIGAR induces a shift from glycolysis to pentose phosphate pathway (PPP), resulting in the regeneration of NADPH, thereby enhancing reductive capacity of the cell^53^. A previous study has shown that knockdown of TIGAR increases the amount of DNA damage caused by epirubicin^54^. It could therefore be that p53-induced upregulation of TIGAR protects cells against ROS-induced DNA damage by increasing flux through the PPP, which is in line with our metabolomics data. But on the other hand, p53 stabilization by Nutlin-3a still mounts protection against DNA damage when cells are deprived of glucose (and hence cannot increase glucose flux through the PPP), arguing that the hormetic response is largely independent of altered glucose metabolism.

Systemic toxicity of chemotherapy is often a limiting factor for cancer treatment. Lowering the doses of chemotherapies to reduce toxicity poses the risk of not eradicating all cancer cells, leaving behind clones that can drive disease relapse. These clones can already be resistant prior to treatment, or acquire resistance due to chemotherapy-induced mutations^55^. In addition, chemotherapy-induced mutations in healthy tissue have been shown to result in secondary cancers^56–58^. p53-induced hormesis as described in this manuscript could potentially be exploited to tackle the toxicity problem in future cancer treatment strategies, taking advantage of the fact that about half of the tumors have lost or impaired p53 function^59^. In a patient with a tumor with impaired p53 function, a p53-inducing drug could confer hormesis and protect the p53 wild-type healthy tissue to chemotherapy, whereas p53 KO cells are not protected. We also noted a slightly increased sensitivity to chemotherapy of p53 KO cells upon Nutlin-3a treatment, which is in line with previous research^60^. The hormetic response in healthy, p53 wildtype cells and increased vulnerability in p53-mutant cells would allow for more efficient tumor killing with less DNA damage-derived systemic side effects. Indeed, we show that Nutlin-3a pretreatment protects p53 WT cells against doxorubicin and etoposide, whereas p53 KO cells remain vulnerable.

MDM2-inhibitors are already being tested in clinical trials as cancer therapy, but with the rationale to (re)activate p53 in p53 wild-type tumors^61^. Unfortunately, many of these trials were ended prematurely due to hematologic, gastrointestinal and other toxicities probably arising from systemic p53 stabilization^62,63^. However, in our proposed approach we envision that p53-stabilization will be used only shortly and at a lower dose as a prophylactic, prior to conventional chemotherapy. Indeed, an intermittent dosing schedule was shown to decrease toxicity of an MDM2 inhibitor^64^. Taken together, the use of MDM2 antagonists to protect healthy tissue might be a promising novel therapeutic approach to reduce toxicity of DNA-damage inducing chemotherapy in healthy tissue. Future work is needed to explore how generalizable this approach would be, using other cell lines, advanced pre-clinical tumor model systems and other chemotherapies with different modes of action.

## Materials and Methods

### Cell culture and treatments

Immortalized retinal pigment epithelial cells (hTERT RPE-1) (ATCC, CRL-4000) were cultured in DMEM-F12, supplemented with 10% FBS (Bodinco), 100 Units Penicillin-Streptomycin (Sigma-Aldrich, P0781) and 2 mM L-glutamine (Sigma-Aldrich, G7513). Cells were cultured at 37 °C and 6% CO_2_. We noted that the exact [D-Ala] that induced lethal DAAO-dependent H_2_O_2_ production at specific timepoints varied between experiments and is likely due to differences in cell density or remaining glucose concentration in the media (PMID: 38548751, PMID: 37392950). We therefore always used a range of D-Ala concentrations to compare conditions. Chemicals that were used are L-alanine (Sigma-Aldrich, A7627), D-alanine (Sigma-Aldrich, A7377), Glucose Oxidase (Sigma-Aldrich, G2133), Nutlin-3 (Cayman Chemicals, 10004372), Hydrogen peroxide (Sigma-Aldrich, H1009), Palbociclib (Selleck Chemicals, S1116), Nocodazole (Sigma-Aldrich, M1404), Etoposide (Sigma-Aldrich, E1383), Doxorubicin (Sigma-Aldrich, D1515). Cells were irradiated using a Gammacell 1000 irradiator.

### Cloning and lentiviral infection

DAAO and HyPer7 and p53 knock-out lines were made as described before^30^. The original plasmid containing GRX1-roGFP2 was a gift from Tobias Dick^65^, and the adapted version for lentiviral delivery with a *Hef1* promoter and a puromycin resistance cassette (backbone pLentiPGK Puro DEST p38KTRClover from Addgene) was kindly provided by Maria Rodriguez-Colman^66^. Lentivirus was made by transfecting HEK293T cells with GRX1-roGFP2 plasmid (localizes throughout the cell) and packaging vectors pRc-cmv-rev1b, pHDM.G and pHDM-Hgpm2. Virus was added to hTERT RPE-1 cells with 8 µg/ml polybrene (Sigma-Aldrich, H9268) and cells were selected with 1 µg/ml puromycin.

### Flow cytometry

For all flow cytometry experiments, medium, wash buffer (PBS) and trypsinized cells were collected on ice. Cells were centrifuged at 1500 RPM for 5 min (Thermo Scientific SL40 centrifuge) to remove media and wash buffer.

For assessment of cell viability by propidium iodide (PI, Sigma-Aldrich, P4170) exclusion, the cell pellet was resuspended in PBS containing 20 µg/ml PI.

For cell cycle profiles, cells were collected as described above, fixed in 70% ethanol and left at 4 °C overnight. On the day of analysis, cells were centrifuged at 1500 RPM for 5 min and resuspended in PBS containing 20 µg/ml PI and 200 µg/ml RNase (Roche, 10109169001).

For BrdU assays, 10 µM BrdU (BD Biosciences, 512420KC) was added to the culture medium 1 hour prior to harvest. Cells were collected and fixed as described above. On the day of analysis, cells were centrifuged at 1500 RPM for 5 min, supernatant was removed and cells were resuspended in 0.1 M HCL solution containing 0.5 mg/ml pepsin (Merck, 1.07197) and left on a tube roller at room temperature for 20 min. Cells were washed with PBS containing 0.1% BSA and 0.5% Tween, cells were centrifuged and pellet was resuspended in 2 M HCL for 12 min at 37 °C to denature the DNA. Borate buffer (0.1 M H_3_BO_3_ set at pH 8.5 with 0.1 M Na_2_B_4_O_7_) was added, cells were centrifuged and pellet was incubated in PBS containing 0.1% BSA, 0.5% Tween and BrdU-FITC antibody (1:50, BD Biosciences, 347583) for 1 h on ice in the dark. Cells were pelleted and resuspended in PBS containing 0.1% BSA, 20 µg/ml PI and 200 µg/ml RNase.

Samples were analyzed on a BD FACS Celesta Cell analyzer (BD biosciences) and analyzed with BD FacsDiva Software. Gating strategies are shown in Supplemantary Fig. 6.

### Live cell imaging

For imaging experiments, cells were seeded in ibiTreated 8-well µ-slides (Ibidi) and imaged on a Cell Observer microscope (Zeiss) using a 10 x magnification and ZEN 2.6 blue software (v.2.6.76.00000). For HyPer7 measurements, cells expressing HyPer7 were imaged using excitation at 385 nm and emission at 475 nm every 10 min. Note: When imaging NES-HyPer7 we always observed an increased HyPer7 ratio at the start of the experiment, regardless of treatment. Therefore, cells were equilibrated in the microscope at least half an hour before adding D-Ala. For FUCCI measurements, cells were imaged every 15 min. For live PI exclusion measurements, medium was replaced with 10 µg/ml PI (Sigma-Aldrich, P4170) at the start of the experiment and cells were imaged every 15 min. The amount of PI positive cells after addition of 0.5% TX-100 (Thermo Scientific, A16046.AP) at the end of the experiment was set as 100%.

### SDS-PAGE and Western Blot

Cells were seeded in 6-well plates and treated with L-Ala or D-Ala. Cells were washed twice with PBS and scraped in 1X sample buffer containing 2% SDS, 5% 2-mercaptoethanol, 10% glycerol, 0.002% bromophenol blue and 300 mM Tris-HCI pH 6.8. Samples were incubated at 95 °C for 10 min and were run on an SDS-PAGE gel. Proteins were transferred on a PVDF membrane. Membranes were blocked for 1 hour at 4 °C in TBS-Tween containing 1% BSA (Sigma-Aldrich, A9647). Membranes were incubated with primary antibody in TBS-Tween overnight at 4 °C and with secondary antibodies in TBS-Tween (1:10000) for 1 hour at 4 °C. Blots were imaged on Image Quant LAS (GE HealthCare) with ImageQuant LAS4000 (version 1.3) software. Contrast was adjusted by linear image processing in Adobe Photoshop.

### Antibodies

Antibodies used for Western Blot were p53 (DO-1) (Santa Cruz, SC-126, lot# E2521, 1:1000), p21 (BD Biosciences, 556430, lot#5073543, 1:5000), TIGAR (Cell Signaling, CS14751S, ref# 02/2015, 1:2500), Tubulin (Millipore, CP06 OS, lot#4079387 1:5000).

### RNA sequencing

Cells were treated with 10 mM D-Ala or L-Ala for 4 hours and sample collection and RNA extraction was performed using the NucleoSpin RNA kit (Macherey-Nagel, 740955) according to the manufacturer’s instructions. RNA-seq libraries were prepared with the Truseq RNA stranded polyA Library Prep Kit (Illumina) according to the manufacturer’s instructions and were sequenced on a NextSeq2000 (Illumina) as 2 × 50 bp paired-end reads. RNA sequencing reads were aligned against human reference genome 38 using Rasflow^67^. The bioconductor package DESeq2 was used to perform differential expression analysis. The bioconductor package fgsea was used to perform gene set enrichment analysis on genes ranked by stat values from DESeq and using hallmark pathways.

### Mass spectrometry

#### LC-MS/MS

Cells (in 6-wells format) were washed with PBS and lysed with 200 ml 8 M Urea in 100mM HEPES (pH 8), after which the lysate was diluted 1.5 fold with 100mM HEPES (pH 8) and 6 mM MgCl2 and 50 U benzonase was added. After a 10-minutes incubation at 37°C, proteins were alkylated in 10 mM Tris(2-carboxyethyl)phosphine hydrochloride (TCEP) and 40mM 2-chloro-acetamide after which proteins were cleaned up with the SP3 protocol^68^. Proteins (∼30 ug) were digested overnight with 0.5ug Trypsin/LysC (Thermo) in 100 mM HEPES (pH 8), 10 mM CaCl2. After acidification with 2% Formic Acid (FA) peptides were separated from the beads and were loaded on C-18 stage tips (Affinisep). After elution from the stage tips, acetonitrile was removed using a SpeedVac and the remaining peptide solution was diluted with buffer A (0.1% FA) before loading. Peptides were separated on a 25 cm in-house made pulled emitter fused silica column (50 µm ID, Polymicro) packed with 1.9 µm aquapur gold C-18 material (dr. Maisch) using a 1 hour gradient (7% to 80% ACN 0.1% FA), delivered by an Vanquish Neo HPLC (Thermo), and electro-sprayed directly into a Orbitrap Astral Mass Spectrometer (Thermo Scientific). The latter was set in data independent mode with a cycle time of 0.6 second for both Faims CV settings (-45V and -65V), in which the full scan was performed at a resolution of 240K and an automatic gain control at 500%. The precursor mass range for both CVs was 380-980 Th in which the peptides, from an isolation window of 2 Th, were fragmented with a normalized collision energy of 25%.

#### Data analysis

Raw files were analyzed with Proteome Discoverer (Thermo)(Version 3.0) with oxidation of methionine set as variable modifications, and carbamidomethylation of cysteine set as fixed modification. The Human protein database of Uniprot (Nov. 2024) was searched with both the peptide as well as the protein false discovery rate set to 1%. Quantfication was done on the MS1 level. The proteomics data will be deposited to the ProteomeXchange Consortium via the PRIDE partner repository (http://www.ebi.ac.uk/pride).

The bioconductor package fgsea was used to perform gene set enrichment analysis on proteins ranked by t-statistic values (log2FC / SE) and using hallmark pathways.

### Metabolomics

#### Materials

Organic solvents were ULC-MS grade and purchased from Biosolve (Boom B.V., Meppel, The Netherlands). Chemicals were analytical grade and purchased from Sigma-Aldrich (Zwijndrecht, The Netherlands). Ultrapure water was obtained fresh from a Milli Q instrument (Merck Millipore, Amsterdam, The Netherlands). A Labconco Centrivap and Thermoshaker were obtained from VWR International (Amsterdam, The Netherlands). A Vanguard UPLC system, an Exploris 480 Orbitrap MS, Compound Discoverer v3.3 software and XCalibur software v4.7 were purchased from Thermo Scientific (Breda, The Netherlands). UPLC columns were obtained from Waters (Etten-Leur, The Netherlands).

#### Liquid Chromatography Mass Spectrometry (LC-MS)

Sample fractions in 1.5 mL Safe-Lock cap Eppendorf tubes were evaporated to dryness in a Centrivap sample concentrator at 40°C. To the dry samples 200 µL methanol (containing 50 µM BHT) and 400 µL chloroform were added. The samples were homogenized by ultrasonification for 5 min and incubated at for 1 h at 30°C and 900 rpm in a thermoshaker. Phase separation was induced by adding 200 µL ultrapure water. The samples were pulse vortex mixed, stored for 10 min at 4°C, and centrifugated for 10 min at 4°C and 17000 x g. Both the upper aqueous phase and lower organic phase were collected in labelled Eppendorf vials and evaporated to dryness in a Centrivap concentrator and stored at -80 until analysis. On the day of analysis the sample residues were dissolved in 100 µL milli Q water and transferred to sample vials kept at 5°C during analysis. Quality control (QC) samples were prepared by pooling 10 µL aliquots from each individual sample. QC samples were analyzed after every tenth experimental sample throughout the sample analysis sequence.

For reversed phase separation either an Atlantis Premier BEH C18 AX UPLC-column (2.1×100 mm, 1.7 μm) for analysis at basic conditions, and an HSS-T3 UPLC-column (2.1×100 mm, 1.8 µm) for analysis at acidic condition. The reversed phase columns were operated at a flow rate of 300 µL min-1 and kept at 30°C. Mobile phase A for analysis at basic conditions consisted of 20 mM ammonium acetate and 5 mM ammonium hydroxide, and at low pH it consisted of 0.1% formic acid. Mobile phase B was acetonitrile in both cases. A step gradient was started upon injection of 5 µL of sample. The gradient steps were 0% B for 1.5 min followed by a 5 min linear increase to 15% B after which the percentage B was increased linearly to 70% during 3.5 min. The gradient was kept at 70% for 2 min, after which the system returned to 0% B in 0.2 min and the column was allowed to re-equilibrate for 7 min prior to a next injection. For lipidomics analysis the residue of the organic phase was dissolved in 100 µL acetonitrile. Analysis was conducted with the system described above using an Acquity BEH C18 column (2.1×100 mm, 1.7 µm) kept at 60°C. The system was operated at a flow rate of 450 μL min^−^1. The mobile phases consisted of 40% acetonitrile also containing 10 mM ammonium acetate (solvent A), and 10% acetonitrile - 90% isopropanol also containing 10 mM ammonium acetate (solvent B) for both negative and positive mode. A 12-min linear gradient of 40–100% B was started after the injection of 5 µL of the sample. The system was kept at 100% B for the next 5 min, after which the system returned to its starting situation. Total runtime was 20 min.

The column outlet was coupled to the Exploris 480 MS. A heated electrospray ionization (HESI) interface was used for ionization. Data were acquired in both negative and positive ionization modes, with spray voltages of 3.6 kV and 2.5 kV, respectively. The capillary temperature was maintained at 320 °C, the auxiliary gas at 11 arbitrary units, the spare gas to 2 arbitrary units, and the S-lens RF level to 50. Spectra were acquired in full-scan mode using the Orbitrap analyzer at a resolution of 180,000, with an automatic gain control of 3 × 106, a maximum injection time of 200 ms, and a mass accuracy of 5 ppm. MS/MS acquisition was performed using a collision energy of 40. For metabolite identification, a QC and standards sample were additionally analyzed by MS/MS. Data analysis was performed using XCalibur Quan and Compound Discoverer 3.3 software (Thermo Scientific), using data from standards and the KEGG, HMDB and lipidmaps databases. QRILC imputation was perfomed using MetaboAnalyst and peak intensity values were normalized to protein concentration as measured by a Pierce BCA Assay Kit (Thermo Fisher, 23225) and subsequently normalized to untreated control.

### Comet assay

Alkaline comet assays were performed using the CometAssay Single Cell Gel Electrophoresis Assay kit (R&D systems 4250-050-K) according to the manufacturer’s instructions. Cells were lysed overnight and DNA staining was performed using Midori Green (Nippon Genetics, MG04) 1:5000 in TE buffer pH7.5 for 30 min at room temperature. Comets were imaged on an EVOS-M5000 microscope (Thermo Fisher). Comets were analyzed using a python script.

### Senescence-associated-β-galactosidase staining

Cells were seeded in 6-well plates, following pretreatment with Nutlin-3a for 4 hours or 24 hours before addition of L-Ala or D-Ala. Subsequently, cells were washed twice with PBS and fixed for 15 min. in 3.7% formalin (37% formalin (Merck, 1.04003) diluted in PBS). Cells were washed twice in PBS and were incubated overnight at 37 °C in SA-β-gal buffer. SA-β-gal buffer consists of 25.3 mM Na2HPO4 (Merck, 1.06580), 7.4 mM Citric Acid (Sigma-Aldrich,1.00241), 150 mM NaCl (Sigma-Aldrich, 31434), 2 mM MgCl2 (Merck, 1.05833), 5 mM Potassium Ferricyanide (Sigma-Aldrich, 702587), 5 mM Potassium Ferrocyanide (Sigma-Aldrich, P9387), this solution was passed through a 0.2 micron filter and 0.1% X-Gal was added (from a 4% X-Gal solution (Meridian Bioscience, BIO37035) in dimethylformamide (Merck 8.22275). Cells were washed twice with PBS, 70% ethanol was added and cells were imaged using the EVOS-M5000 microscope (Thermo Fisher). Relative amount of SA-β-GAL positive cells represents the percentage of the area that is positive for the staining.

### Measuring DAAO activity by oxygen consumption rate

DAAO activity was assessed by measuring oxygen consumption rate (OCR) as described before^32^. OCR was measured using a Seahorse XFe24 analyzer (Agilent) with Wave (version 2.6.1.56) software was used to measure OCR. 24-well V7 Seahorse culture plates (Agilent, 100777) were coated with 50 µl of 50 µg/ml rat tail collagen (Corning, 354236) in 0.1% acetic acid and washed with PBS. Cells were seeded, leaving 4 wells empty for background correction. On the day of the assay, medium was replaced with XF DMEM (Agilent, 103575) supplemented with 2 mM L-Glutamine (Sigma-Aldrich, G7513), 17.5 mM glucose (Gibco, A2494001), 1 mM Na-Pyruvate (Lonza, BE13-115E). Oligomycin A (final concentration 2 µM, Cayman Chemical, 11342) and L/D-Alanine was added to injection ports.

## Data availability

Raw sequencing reads will be available in SRA. Count table of RNA-seq data will be deposited in GEO. Proteomics data will be uploaded via the ProteomeXchange Consortium via the PRIDE partner repository. Metabolomics data will be uploaded on MetaboLights.

## Acknowledgements

We would like to thank Livio Kleij for technical support with live imaging and our colleagues at the Center for Molecular Medicine (UMC Utrecht) and the Van Boxtel lab (Princess Maxima Center) for their valuable input and suggestions. This work was funded by a grant from the Dutch Cancer Society (KWF Kankerbestrijding, project number 14798) to TBD. BMTB and RvB are members of the Oncode Institute, which is partly funded by the Dutch Cancer Society (KWF Kankerbestrijding).

## Author contributions

J.P.K., P.E.P., and T.B.D. designed, performed and analysed experiments. W.T.F.d.T. P.S.A., R.v.E., S.E.B and H.R.V performed proteomic analysis. M.C.G and E.C.A.S performed metabolomic analysis. D.M.G performed mapping of mRNA sequencing data. R.K. made a script to automatically analyze comet assays. W.T.F.d.T., B.M.T.B. and R.v.B. provided critical input on the experimental setup, results and the manuscript. J.P.K. and T.B.D. conceived the study and wrote the manuscript.

