## Supplementary Figures 1-6 for "Hydrogen Peroxide induces resistance to DNA damage in a localization and p53 dependent manner"

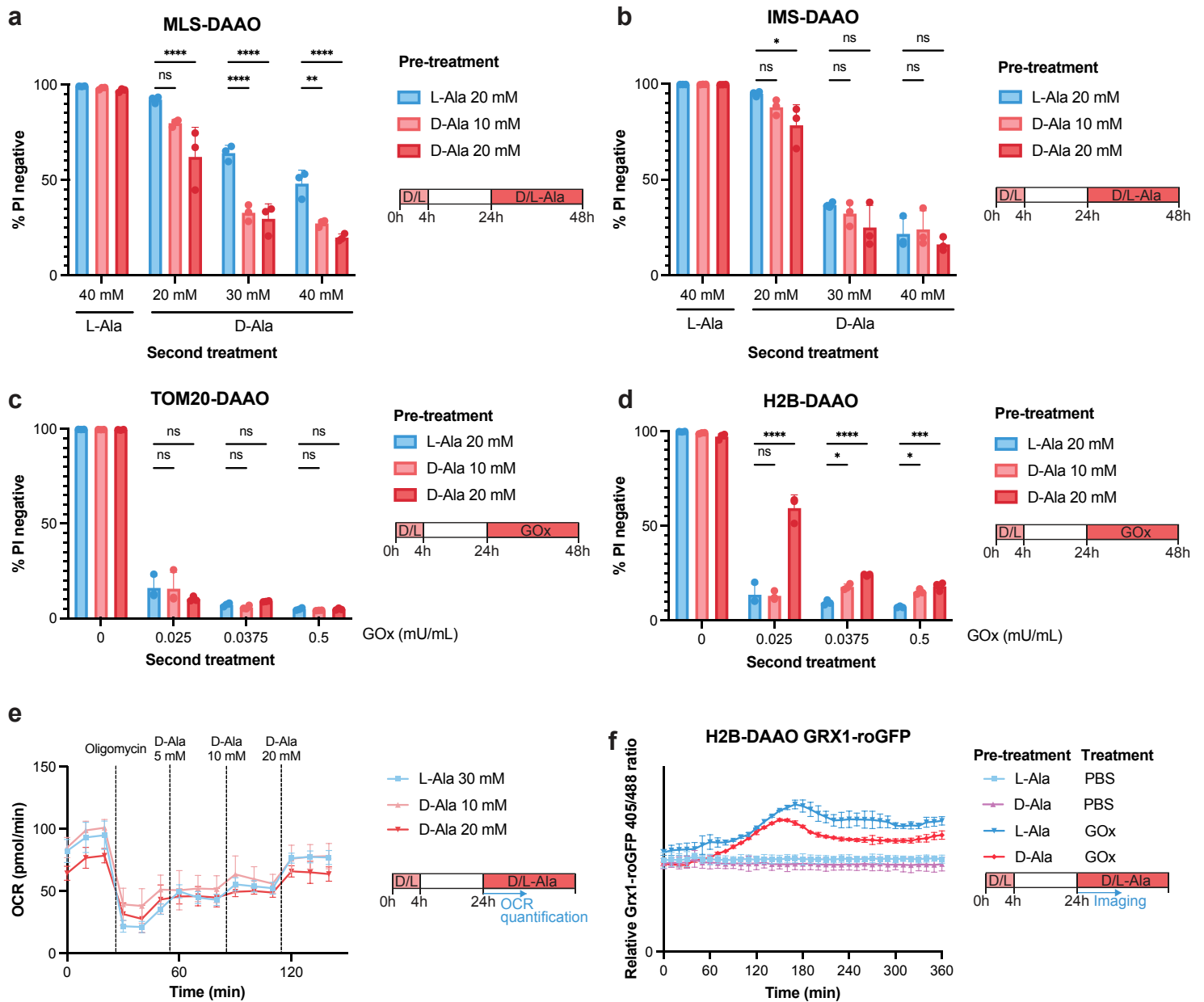

### Supplementary Fig. 1

**a-b** Quantification of cell viability by PI exclusion of RPE1-hTERT MLS-DAAO cells (**a**) and RPE1-hTERT IMS-DAAO cells (**b**) that were pretreated with L-Ala or D-Ala for 4 hours, followed by 20 hours of recovery and treatment with L-Ala or D-Ala. Dots represent 3 biological replicates of ~10.000 cells each. Data represents mean ± SD. Two-way Anova with Bonferroni correction was performed (ns $p > 0.05$, \*$p \leq 0.05$, \*\*$p \leq 0.01$, \*\*\*\*$p \leq 0.0001$).

**c-d** Quantification of cell viability by PI exclusion of RPE1-hTERT TOM20-DAAO cells (**c**) and RPE1-hTERT H2B-DAAO cells (**d**) that were pretreated with L-Ala or D-Ala for 4 hours, followed by 20 hours of recovery and treatment with Glucose Oxidase (GOx). Dots represent 3 biological replicates of ~10.000 cells each. Data represents mean ± SD. Two-way Anova with Bonferroni correction was performed (ns $p > 0.05$, \*$p \leq 0.05$, \*\*\*$p \leq 0.001$, \*\*\*\*$p \leq 0.0001$).

**e** Oxygen consumption rate (OCR) measurements as a measure for enzymatic activity of H2B-DAAO upon addition of D-Ala. Cells were pretreated with L-Ala or D-Ala for 4 hours, followed by 20 hours of recovery. Subsequently, DAAO activity was measured by changes in OCR upon D-Ala titration. Data represents mean ± SD of 3 biological replicates.

**f** Relative GRX1-roGFP2 405/488 ratio of RPE1-hTERT H2B-DAAO cells expressing GRX1-roGFP2 that were pretreated with 20 mM L-Ala or D-Ala for 4 hours followed by 20 hours of recovery. Subsequently, cells were imaged and 0.125 mU/mL Glucose Oxidase (GOx) was added. Data represents mean ± SD of 2 biological replicates.

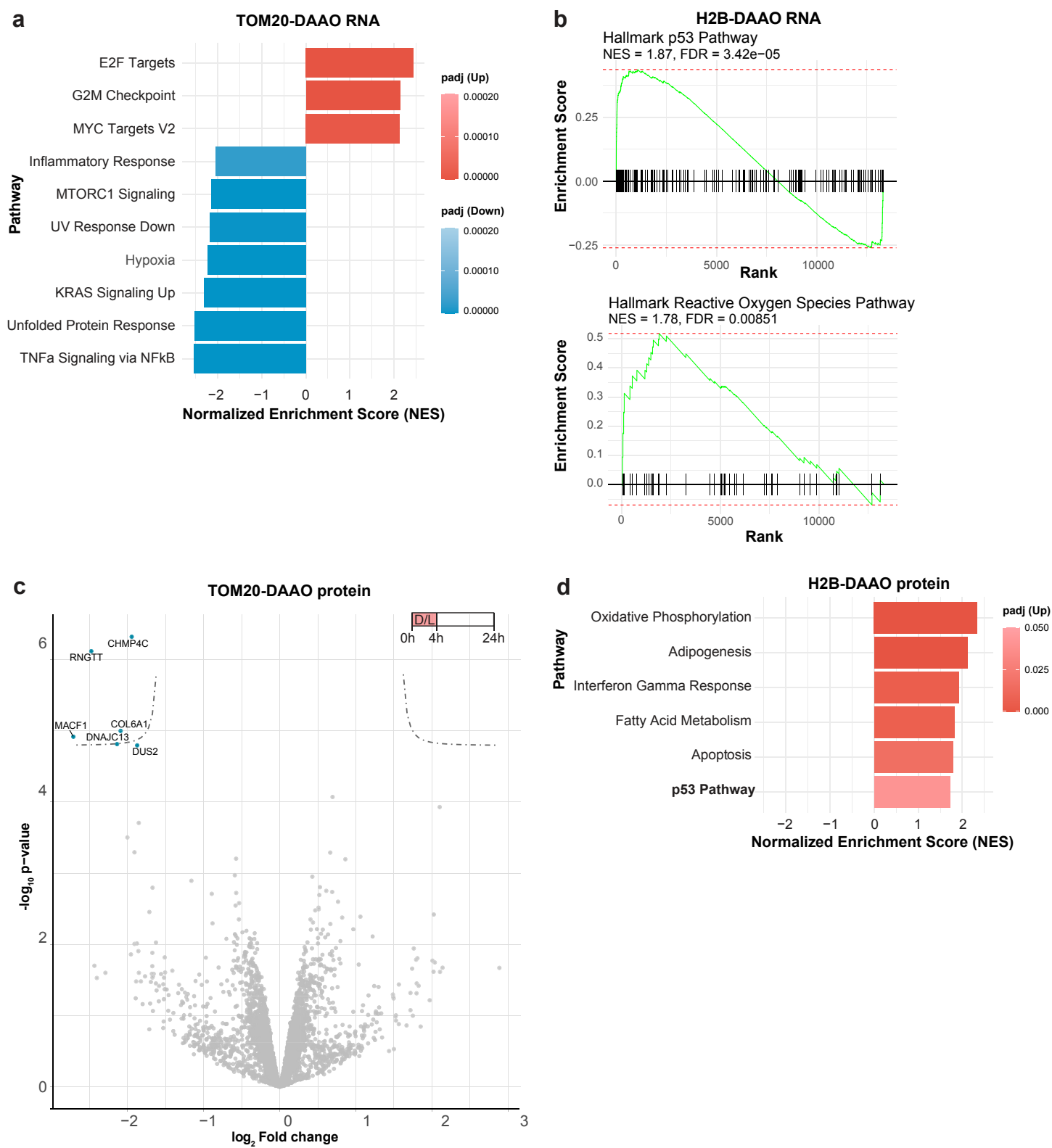

### Supplementary Fig. 2

- **a** Top 10 Hallmark pathways that are changed upon D-Ala treatment in RPE1-hTERT TOM20-DAAO cells, determined using gene set enrichment analysis (GSEA) of mRNA sequencing data.
- **b** Gene set enrichment analysis plots of hallmark pathways (p53 pathway and Reactive Oxygen Species Pathway) enriched in mRNA sequencing data of RPE1-hTERT H2B-DAAO cells upon D-Ala treatment.
- **c** Volcano plot depicting differentially expressed proteins comparing L-Ala versus D-Ala treatment of 20 mM for 4 hours, followed by 20 hours of recovery in RPE1-hTERT TOM-DAAO cells. Blue dots represent significantly upregulated or downregulated proteins (FDR≤0.05).

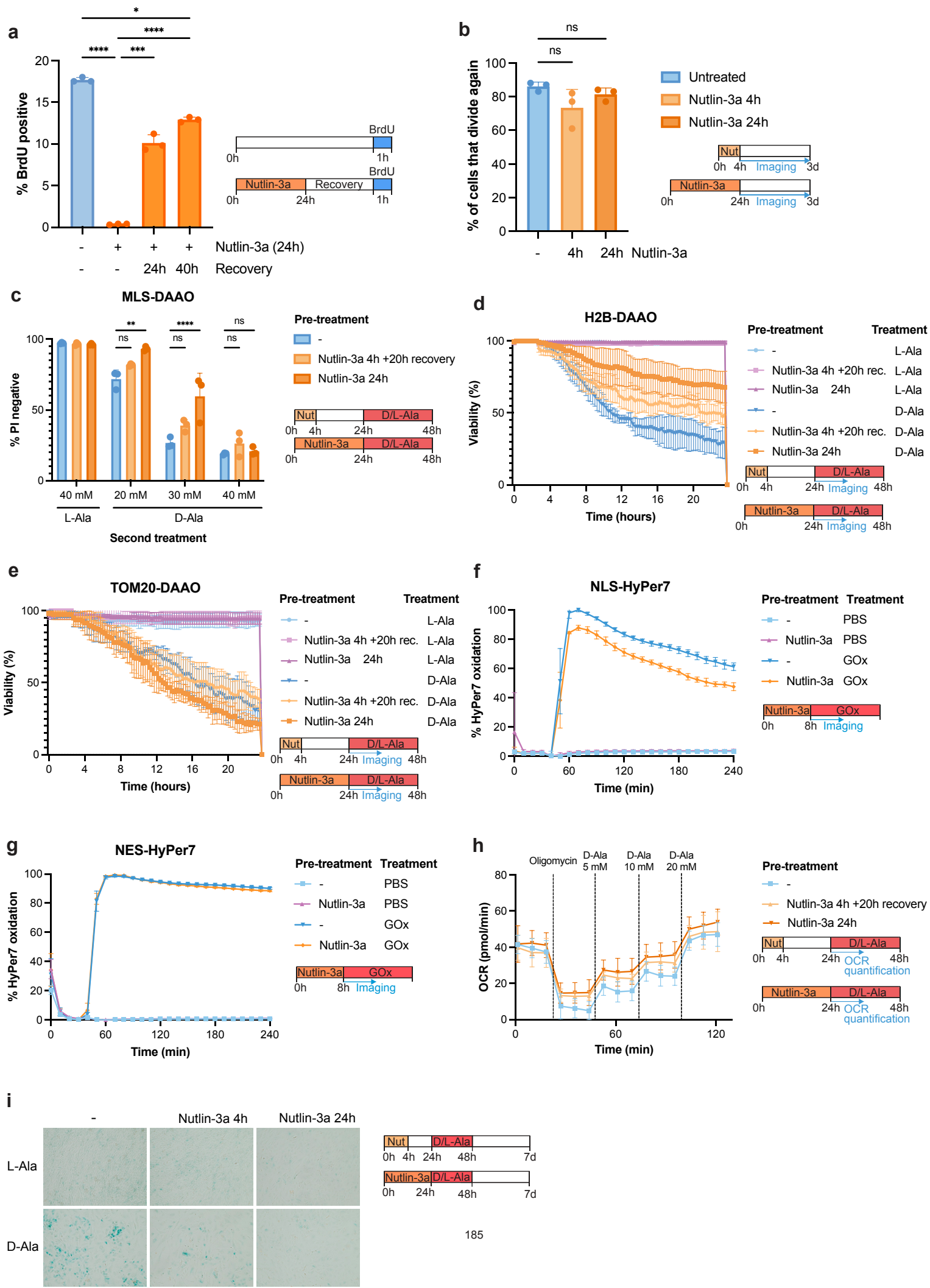

#### Supplementary Fig. 3

**a** Quantification of BrdU incorporation assay by flow cytometry. BrdU was added for the last hour of the experiment. Dots represent 3 biological replicates of ~10.000 cells each. Data represents mean +/- SD. Two-way Anova with Bonferroni correction was performed (ns $p > 0.05$, $p \leq 0.05$, \*\*\*$p \leq 0.001$, \*\*\*\*$p \leq 0.0001$).

**b** Quantification of live imaging of cells expressing the FUCCI marker to indicate cell cycle phase. Cells were treated with Nutlin-3a for 4 or 24 hours and during recovery, the amount of cells that started dividing again was counted. Dots represent 3 biological replicates of ~30 cells each. Data represents mean +/- SD. Two-way Anova with Bonferroni correction was performed (ns $p > 0.05$).

**c** Quantification of cell viability by PI exclusion of RPE1-hTERT MLS-DAAO cells that were pretreated with Nutlin-3a (10 μM) for 4 hours and 20 hours of recovery or 24 hours, followed by treatment with L-Ala or D-Ala. Dots represent 3 biological replicates of ~10.000 cells each. Data represents mean +/- SD. Two-way Anova with Bonferroni correction was performed (ns $p > 0.05$, \*\*$p \leq 0.01$, \*\*\*\*$p \leq 0.0001$).

**d-e** Quantification of cell viability of RPE1-hTERT H2B-DAAO cells (d) or RPE1-hTERT TOM20-DAAO cells (e) by measuring PI positive cells during live imaging. The amount of PI positive cells was normalized to the total amount of cells in the well at the end of the experiment after lysing the cells using 10% Triton X-100. Cells were pretreated with Nutlin-3a (10 μM) for 4 hours and 20 hours of recovery or 24 hours, followed by treatment with L-Ala or D-Ala.

**f-g** HyPer7 measurements of RPE1-hTERT H2B-DAAO cells with nuclear NLS-HyPer7 (f) or cytosolic NES-HyPer7 (g) that were pretreated with Nutlin-3a (10 μM) for 8 hours. Subsequently, cells were imaged and 0.25 mU/mL Glucose Oxidase (GOx) was added. Data represents mean +/- SD of 4 biological replicates.

**h** Oxygen consumption rate (OCR) measurements as a measure for enzymatic activity of H2B-DAAO upon addition of D-Ala. Cells were pretreated with Nutlin-3a (10 μM) for 24 hours. Subsequently, DAAO activity was measured by measuring changes in OCR. Data represents mean +/- SD of 3 biological replicates.

**i** Representative images of Senescence associated-β-galactosidase (SA-β-gal) staining of RPE1-hTERT H2B-DAAO cells that were pretreated with Nutlin-3a (10 μM) for 4 hours and 20 hours of recovery or 24 hours, followed by treatment with 10 mM L-Ala or D-Ala for 48 hours. Subsequently, cells were kept in culture for 4 more days to allow for expression of SA-β-gal. Quantification is shown in Fig. 3f.

**a**

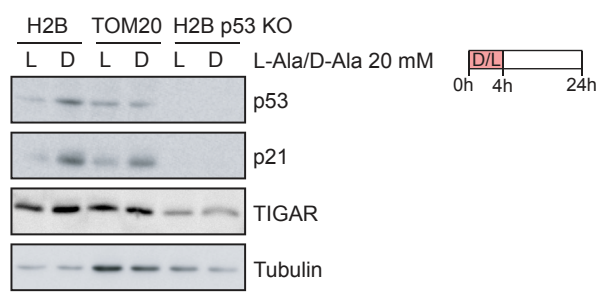

#### Supplementary Fig. 4

**a** Western Blot of H2B-DAAO cells, TOM20-DAAO cells and H2B-DAAO p53 KO cells that were treated for 4 hours with 20 mM L-Ala or D-Ala, followed by 20 hours of recovery.

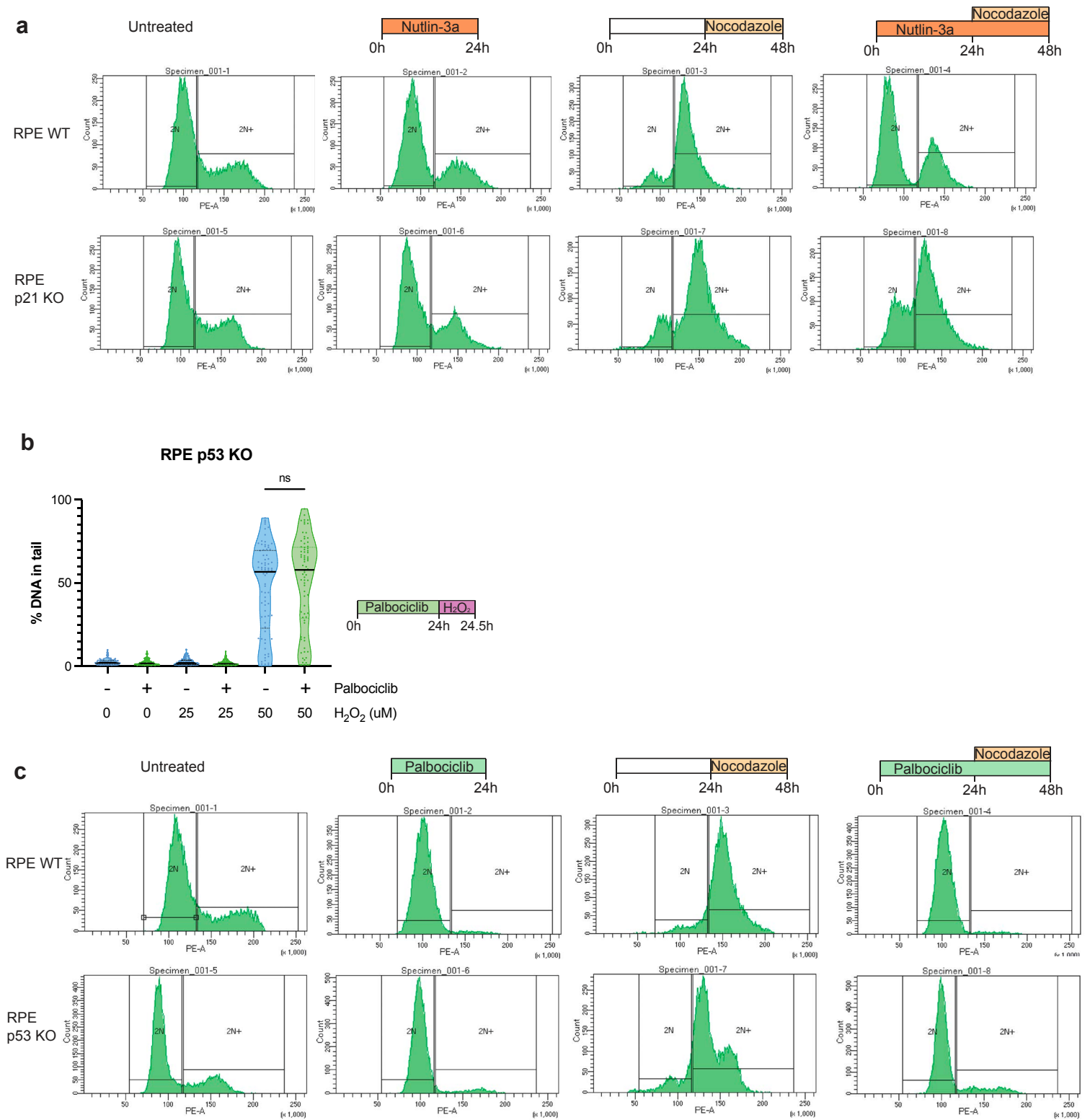

### Supplementary Fig. 5

**a** Cell cycle profiles to confirm Nutlin-3a-induced arrest of p21 WT cells and no arrest in and p21 KO cells. Cells were stained using PI and ~10,000 cells per condition were measured.

**b** Quantification of alkaline comet assay of RPE1-hTERT p53 KO cells pretreated with palbociclib for 24 hours, followed by 30 min of H<sub>2</sub>O<sub>2</sub> treatment. Line represents mean of ~70 comets per condition. Kruskal-Wallis test with Dunn's post-hoc test was performed (ns p > 0.05).

**c** Cell cycle profiles to confirm G1 arrest after palbociclib treatment of RPE1-hTERT p53 WT and p53 KO cells. Cells were stained using PI and ~10,000 cells per condition were measured.

#### a Propidium iodide exclusion

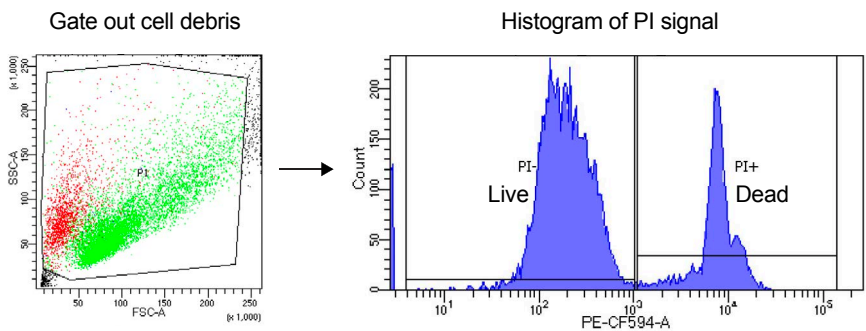

#### b BrdU incorporation

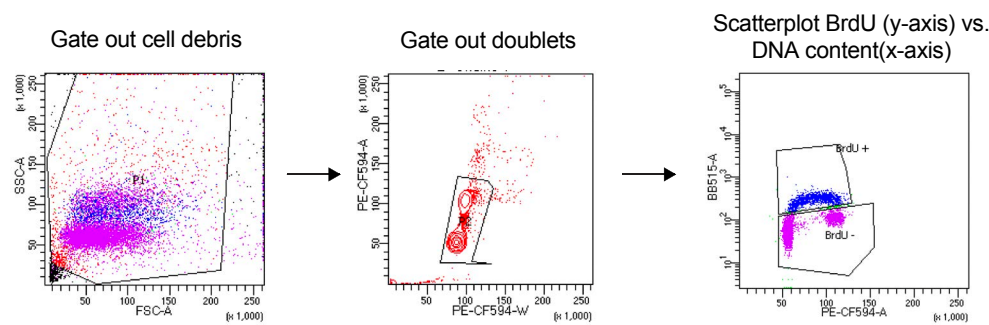

#### c Cell cycle profile

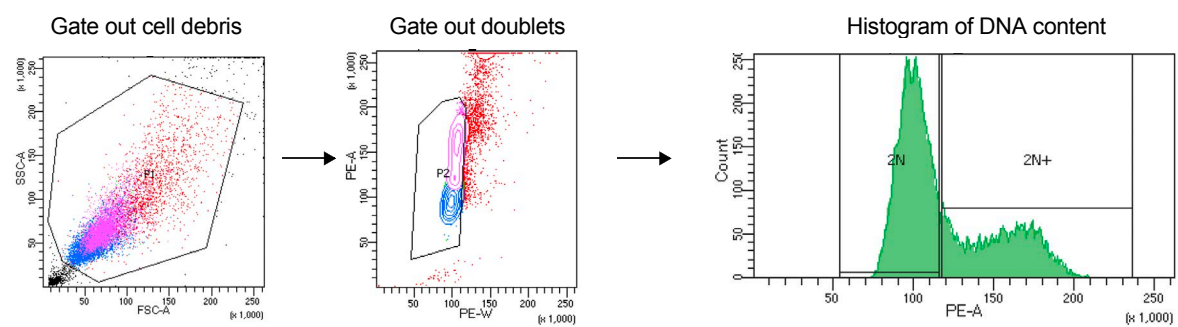

### Supplementary Fig. 6

**a** Gating strategy to determine cell viability by PI exclusion. Cellular debris is gated out using the forward scatter (FSC) and side scatter (SSC). PI positive and negative cells are quantified in the histogram of the PI staining.

**b** Gating strategy to determine cell proliferation by BrdU incorporation. Cellular debris is gated out using the forward scatter (FSC) and side scatter (SSC). Doublets are gated out using PI staining (PE-CF594-A/W). BrdU positive and negative cells are quantified in a plot depicting the BrdU staining (y-axis) and PI staining (x-axis).

**c** Gating strategy to determine cell cycle profiles using a PI staining. Cellular debris is gated out using the forward scatter (FSC) and side scatter (SSC). 2N and 2N+ cells are quantified in the histogram of the PI staining.
